# ProtaBank: A repository for protein design and engineering data

**DOI:** 10.1101/272211

**Authors:** Connie Y. Wang, Paul M. Chang, Marie L. Ary, Benjamin D. Allen, Roberto A. Chica, Stephen L. Mayo, Barry D. Olafson

## Abstract

We present ProtaBank, a repository for storing, querying, analyzing, and sharing protein design and engineering data in an actively maintained and updated database. ProtaBank provides a format to describe and compare all types of protein mutational data, spanning a wide range of properties and techniques. It features a user-friendly web interface and programming layer that streamlines data deposition and allows for batch input and queries. The database schema design incorporates a standard format for reporting protein sequences and experimental data that facilitates comparison of results across different data sets. A suite of analysis and visualization tools are provided to facilitate discovery, to guide future designs, and to benchmark and train new predictive tools and algorithms. ProtaBank will provide a valuable resource to the protein engineering community by storing and safeguarding newly generated data, allowing for fast searching and identification of relevant data from the existing literature, and exploring correlations between disparate data sets. ProtaBank invites researchers to contribute data to the database to make it accessible for search and analysis. ProtaBank is available at https://protabank.org.

**Impact:** The ProtaBank database provides a central repository for researchers to store, query, analyze, and share all types of protein engineering data. This modern database will serve a pivotal role in organizing protein engineering data and leveraging the increasingly large amounts of mutational data being generated. Together with the analysis tools, it will help scientists gain insights into sequence-function relationships, support the development of new predictive tools and algorithms, and facilitate future protein engineering efforts.

**Abbreviations:**

3D: three-dimensional
API: application programming interface
AWS: Amazon Web Services
BLAST: Basic Local Alignment Search Tool
*C*_m_: concentration of denaturant at midpoint of unfolding transition
CSV: comma-separated values
Δ*G*: Gibbs free energy of folding/unfolding
Gβ1: β1 domain of Streptococcal protein G
GdmCl: guanidinium chloride
*k*_cat_: catalytic rate constant
*K*_d_: dissociation constant
MIC: minimum inhibitory concentration
PDB: Protein Data Bank
PE: protein engineering
RDS: Relational Database Services
REST: Representation State Transfer
*T*_m_: melting temperature

## Introduction

Recent advances in gene synthesis, microfluidics, deep sequencing, and microarray techniques have greatly facilitated the ability of researchers to construct and assess large libraries of variant protein sequences.^1–4^ Thousands or even millions of sequence variants can now be generated and screened in an ultrahigh-throughput fashion. This rapid generation of large sets of mutational data has enabled comprehensive mappings between protein sequence and function for properties such as stability, binding affinity, and catalytic activity.^5–7^ Deep mutational scanning approaches have been used to study protein fitness landscapes, discover new functional sites, and engineer proteins with new and improved properties.^8, 9^ Many groups are now using these techniques to generate large amounts of protein engineering (PE) data—a trend that is expected to grow in the future.

The field of PE thus appears to be entering into a state reminiscent of the early days of widespread structure determination and genome sequencing. The Protein Data Bank^10, 11^ (PDB) (www.rcsb.org) and GenBank^12^ were created because scientists recognized the importance of organizing the vast number of protein structures and nucleic acid sequences into databases with standardized formats. Since their inception, these open-access databases have grown exponentially and have proven to be extremely valuable resources for the scientific community. A similar situation now exists with the rapid expansion in PE data. Unfortunately, there is no central repository to store all the PE data being generated, no standardized format for describing it, and no simple means of sharing the data with collaborators.

Here, we present ProtaBank, a database for storing, querying, analyzing, and sharing PE data. This type of information (mutant protein sequences and their associated experimental assay data) is not stored in GenBank or the PDB, and although other databases store some of the data types included in ProtaBank, they are not designed for all types of PE data or have limited tools for inclusion of large amounts of mutant information. Of these, the best known are ProTherm^13^ (thermodynamic database for wild-type and mutant proteins), UniProt^14^ (protein sequences with annotations), and BRENDA^15^ (enzymes and metabolic information). ProTherm is limited to thermodynamic protein stability data and has not been updated since 2013. UniProt and BRENDA were not designed for PE data, and although some mutant data is included, storage and retrieval is limited. The Protein Mutant Database^16^ includes PE data on a broad range of protein properties, but has not been updated in over a decade. Recently, a number of mutant databases were developed to facilitate the study of protein-protein interactions. These include SKEMPI^17^ and PROXiMATE,^18^ which contain thermodynamic data for mutant protein-protein complexes, and AB-Bind,^19^ which focuses on binding data for a select set of antibody-antigen complexes. Overall, the mutational data that is available tends to be scattered across many different specialized databases.

ProtaBank provides a single repository for all types of PE data, spanning a wide range of properties, including those related to activity, binding, stability, folding, and solubility. The database accommodates mutational data obtained from diverse approaches, including computational and other types of rational design, saturation mutagenesis, directed evolution, and deep mutational scanning. Unlike many other mutant databases, ProtaBank stores the entire protein sequence for each of the variants instead of just the mutations and provides detailed descriptions of the experimental assays used. These features are incorporated to allow for accurate comparisons of measurements across multiple studies or groups, making it easier to identify trends and determine how different assays, parameters, or conditions affect the results.

We stress the importance of a standardized format for reporting PE data that allows accurate comparisons between different data sets, and anticipate that the ProtaBank format will become an industry-wide standard used by the entire PE community. This will facilitate sharing PE data with collaborators and will improve the usability of PE datasets for data mining and other analysis methods. The ProtaBank database, together with its analysis and visualization tools, will help scientists gain insights into sequence-activity and structure-activity relationships, improve our understanding of how proteins function, and ultimately facilitate the design of proteins with new and improved properties. The database should also accelerate the development of new predictive tools and algorithms, and lead to improved methods for computational protein design and engineering.

## Database Construction and Content

### Overview

ProtaBank has three main components: (1) a web interface and application programming interface (API) for data deposition, (2) a back-end relational database for data storage, and (3) a web interface and API for data searching and analysis. The design and workflow for ProtaBank is summarized in Figure 1. Users can submit data into the database through the web interface; access to external databases such as PubMed,^20^ the PDB, and UniProt are provided to facilitate the entry of publication information, structural data, and sequence data. In addition, a Representation State Transfer (REST) API layer is provided for batch submission of data. All data undergoes validation and curation before final submission into the database. One can use the web interface or the REST API to search and filter studies and data based on PubMed ID, PDB ID, UniProt accession number, protein name, protein sequence, assay, or publication information. More advanced analysis and comparison tools are also available via the web interface. For example, users can do a sequence search with the Basic Local Alignment Search Tool (BLAST)^21^ and use visualization tools to map mutational data onto a PDB structure.

**Figure 1.**
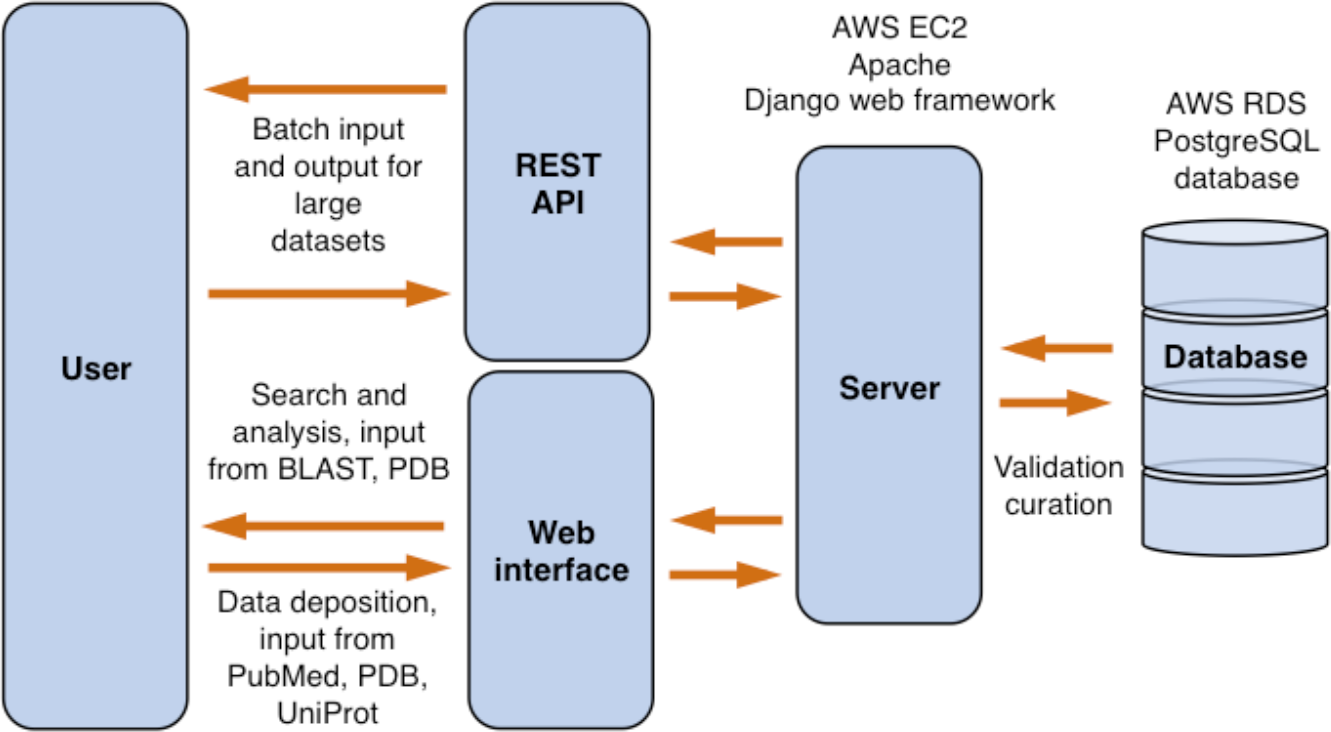
ProtaBank design. Users can interact with ProtaBank through the web interface or the REST API. Data sent to the server is validated and curated before final submission into the database. AWS, Amazon Web Services; RDS, Relational Database Services.

### Database schema

ProtaBank is implemented as a relational database using the PostgreSQL database. The highest level of organization is a study corresponding to a PE effort. Each study has four core tables to describe the PE data: sequence_complex, assay_expassay, data_expfdatum, and data_units which respectively represent the sequence of a given protein mutant, the experimental assay that was used to probe the property of interest, the numerical value obtained for the mutant (i.e., the assay results), and the units associated with the numerical value (Fig. 2). ProtaBank also has separate corresponding tables to represent computational protocols and derived quantities, and to store qualitative data (e.g., folded/unfolded) or data expressed in terms of a range or limit (e.g., 20-30, >100). In addition to the core data tables, each study includes publication information, structural data on the protein that was engineered (i.e., the PDB file, if available), and experimental gene construct information. This information adds context and additional query and filter parameters to the PE data. Non-published PE studies can also be input in a similar fashion. In these cases, the researchers and organizations involved are specified instead of the authors and affiliations. Depositors of non-published results may embargo the release of the data until publication.

**Figure 2.**
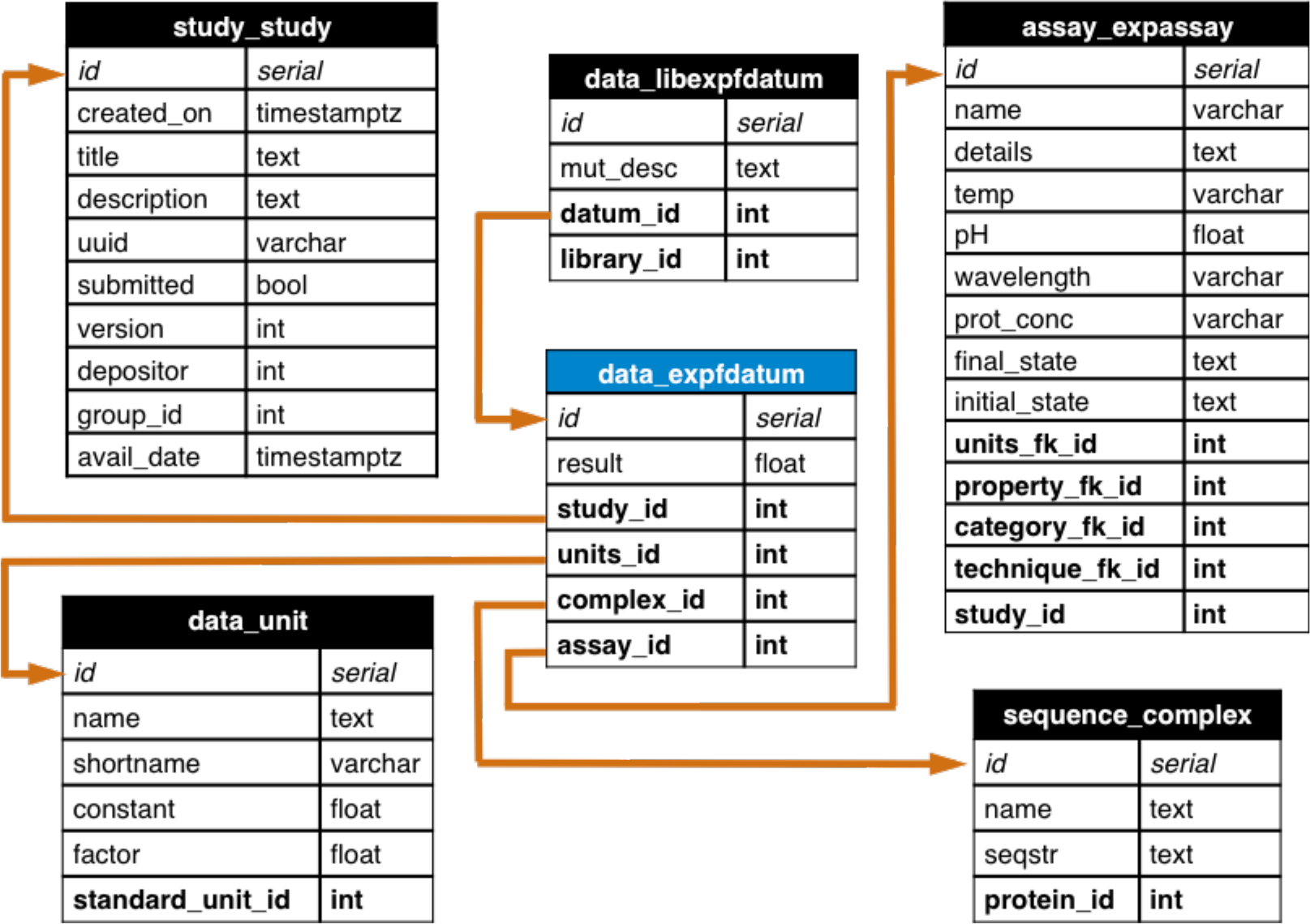
ProtaBank database schema showing the table for experimental data represented by a number (data_expfdatum, blue header) and all tables with foreign key relationships to it. Each table shows the field name (left) and the variable type for the field (right). Each datum in the data_expfdatum table has a foreign key relationship (orange arrow) to a study table (study_study) that organizes the context in which the experiments were performed, an assay table (assay_expassay) that describes the procedure used to obtain the measurement, a sequence table (sequence_complex) that holds the protein sequence of the mutant, and a units table (data_unit) that describes the units of the result. For data that is part of a mutant library, a foreign key links it to the library table (data_libexpfdatum). Analogous tables exist for data obtained from computations/simulations, derived data, and qualitative or range data. uuid, universally unique ID; avail_date, date study is available for public; seqstr, sequence string; temp, temperature; prot_conc, protein concentration; timestamptz, time stamp with time zone; varchar, variable character; bool, boolean; int, integer; float, floating point number; fk, foreign key.

The ProtaBank schema design incorporates two crucial elements: (1) the full amino acid sequence of the protein is stored to facilitate comparison of mutants across different assays and studies, and (2) for each assay, information about the protein property measured, the assay conditions and techniques used, and the units of the resulting data is collected in addition to the results. Although these requirements necessitate more upfront effort in deposition and curation, we believe they are necessary to enable useful comparisons of results across different studies. Our reasoning is as follows.

First, PE studies and databases^13,16^ typically describe a mutant by listing the changes to its protein sequence relative to a specified starting sequence. However, the starting sequences used in engineering a given protein are often not the same across studies, which can cause confusion and makes comparisons challenging. The wild-type protein is not always used; residues may be changed, added, or deleted at the termini, for example, to facilitate expression or purification, or substitutions may be made to make the protein more amenable to the assay conditions. Many mutant databases only store the mutational data for the positions mutated. For example, M3A+V5L+S19T might be used to identify a mutant that has been mutated to Ala, Leu, and Thr at positions 3, 5, and 19, respectively; the rest of the sequence (the background in which the mutations were made) is either not given or not recorded. Not knowing the entire sequence for each mutant confounds comparisons, as any differences in the reported results could be due to differences in the background residues. ProtaBank addresses this issue by providing web forms and an API that parses the input mutant information to return the full sequence so it can be stored as such, allowing for a straightforward comparison of mutants across studies. This feature also makes it possible to validate the accuracy of mutant data provided in the WT#MUT (wild-type amino acid, residue #, mutant amino acid) format; i.e., the wild-type amino acid listed for each of the mutated positions is compared to what is specified in the starting sequence, and any discrepancies are flagged.

Second, comparison across studies may be difficult due to differences in assay conditions or techniques, which can greatly affect the results.^22–24^ The ProtaBank schema takes these issues into account. As outlined above, the database uses the assay_expassay table to describe the procedure that was used to determine a given protein property. This table has foreign key relationships with a series of other tables (category, property, technique, units) that help categorize and describe the many ways these properties can be measured. The category table provides the general type of protein property that was engineered or studied (e.g., stability, activity, binding). The property table is more specific and describes the property that was actually measured and gave rise to the result [e.g., melting temperature (*T*_m_), catalytic rate constant (*k*_cat_), dissociation constant (*K*_d_)]. The categories and properties currently included in ProtaBank are listed in Supporting Information, Table S1. Commonly used experimental or computational techniques are also provided to indicate how the property was assayed (e.g., circular dichroism, surface plasmon resonance). Note that the properties and techniques supplied are not comprehensive, and users can enter additional ones. Finally, the units table contains commonly used units that are appropriate to the property measured. For example, the units available for the Gibbs free energy of folding/unfolding (Δ*G*) are kcal/mol and kJ/mol. This level of description is designed to provide enough detail so that data collected from different sources can be compared and analyzed appropriately.

### Data deposition and curation

ProtaBank will serve as a curated and continuously updated repository for PE studies. Thus far, data in ProtaBank has come directly from the published literature and has been manually entered by ProtaBank developers. To aid in the future data collection process, ProtaBank is designed to accept data input directly from the researchers who performed the study, a strategy that has proved effective in populating other biological databases such as the PDB, GenBank, ArrayExpress,^25^ and WormBase.^26^ Any user can input a study via the ProtaBank data submission tools. Two modes of data deposition are provided: an interactive web interface that supports upload of data in a spreadsheet format [i.e., via a comma-separated values (CSV) file] (see Supporting Information, Fig. S1), and a REST API layer that allows for programmatic batch upload of data (see https://protabank.org/docs for details).

ProtaBank data deposition tools are designed to accept the wide range of data generated in PE efforts and to automate the process so as to facilitate entry and ensure accuracy. Publication details (e.g., authors, title, journal, date, abstract) can be fetched from PubMed, and the protein sequence can be retrieved from the PDB or UniProt. If available, structural data for the protein can be fetched from the PDB.

The database schema requires a description of the methods used to assay the protein mutants. For each assay, one must specify an assay name, the general category of protein property that was engineered or studied, the specific property measured, the technique employed, and the units used. All items except the assay name are specified by selecting from options in a drop-down menu. Additional details can be included if desired. By entering this information, assays can be clearly defined and compared.

PE data can be input in two forms: as individual sequences or as a mutant library (a set of mutant sequences obtained by mutating a specified set of residues in a protein). Mutational data can be uploaded from a CSV file or it can be entered manually on the web form. The data entry page for a mutant library is shown in Supporting Information, Figure S1. To specify a mutant library, the user must first enter the starting protein sequence from which the mutations were made. All mutants in the library can then be described either by their full sequence or as mutations from this starting sequence. Two formats are allowed for the latter: (1) the WT#MUT format (e.g., M3A+V5L+S19T) and (2) the mutated residue range/list format in which a range or list of residues is specified that correlates positions in the starting sequence with the amino acids given in the CSV file (e.g., QRS for residues 3-5; or QRS for residues 3,5,7). ProtaBank then takes the description of mutants entered, parses the data, and stores it as full amino acid sequences.

All submitted data is validated to ensure data integrity before inclusion in the database. Automated tests are first performed to ensure that: (1) the data falls within the correct range of values (e.g., temperature in K must be a non-negative number), (2) the assigned units are appropriate for the assayed property, and (3) the amino acid listed for wild type is consistent with that specified in the starting sequence (for mutants described in the WT#MUT format). Outliers in a data set are also flagged and the submitter is asked to check for accuracy. Currently, ProtaBank developers also check studies manually for sequence and data accuracy, appropriate specification of protein properties, and proper description of assays. Potential errors are sent back to the submitter for review.

### Search and analysis tools

ProtaBank provides several search and analysis tools that allow users to: (1) browse and search for relevant studies queried by publication/study details (title, abstract, author), protein name, PDB ID, UniProt accession number, or protein sequence, (2) identify data and mutants related to a given protein sequence by BLAST search, (3) visualize mutational data mapped onto a three-dimensional (3D) protein structure, and (4) compare and correlate data measured using different assays. Figure 3(A) shows a screenshot of the web interface in which the “Browse all submitted studies” tool was used to filter studies by protein name (“ubiquitin”). Figure 3(B) shows a screenshot in which study analysis tools were used to visualize mutational data on a 3D protein structure. The visualizer is based on PV, an open-source javascript protein viewer (https://biasmv.github.io/pv/index.html) that was extended to allow mutations to be represented on the 3D structure using different color schemes. These include coloring by secondary structure, gradient, minimum, maximum, median, mean, proportion above a reference value, and median deviation from a reference value. In the study depicted here, Jacquier *et al.* investigated the effects of mutations on TEM-1 β-lactamase activity by computing the amoxicillin minimum inhibitory concentration (MIC) score for ~990 point mutants.^27^ Figure 3(B) shows the crystal structure of TEM-1 (PDB ID: 1BTL)^28^ displayed with the backbone colored by the MIC score. In this case, the median deviation from the wild-type value is shown, with residues colored from red to white to blue depending on whether the value is less (red) or greater (blue) than the wild-type value at that position, with white representing an equal value. Pointing the cursor at a residue (e.g., Leu57) highlights it in yellow and displays additional information for that residue in the tables below and to the right.

**Figure 3.**
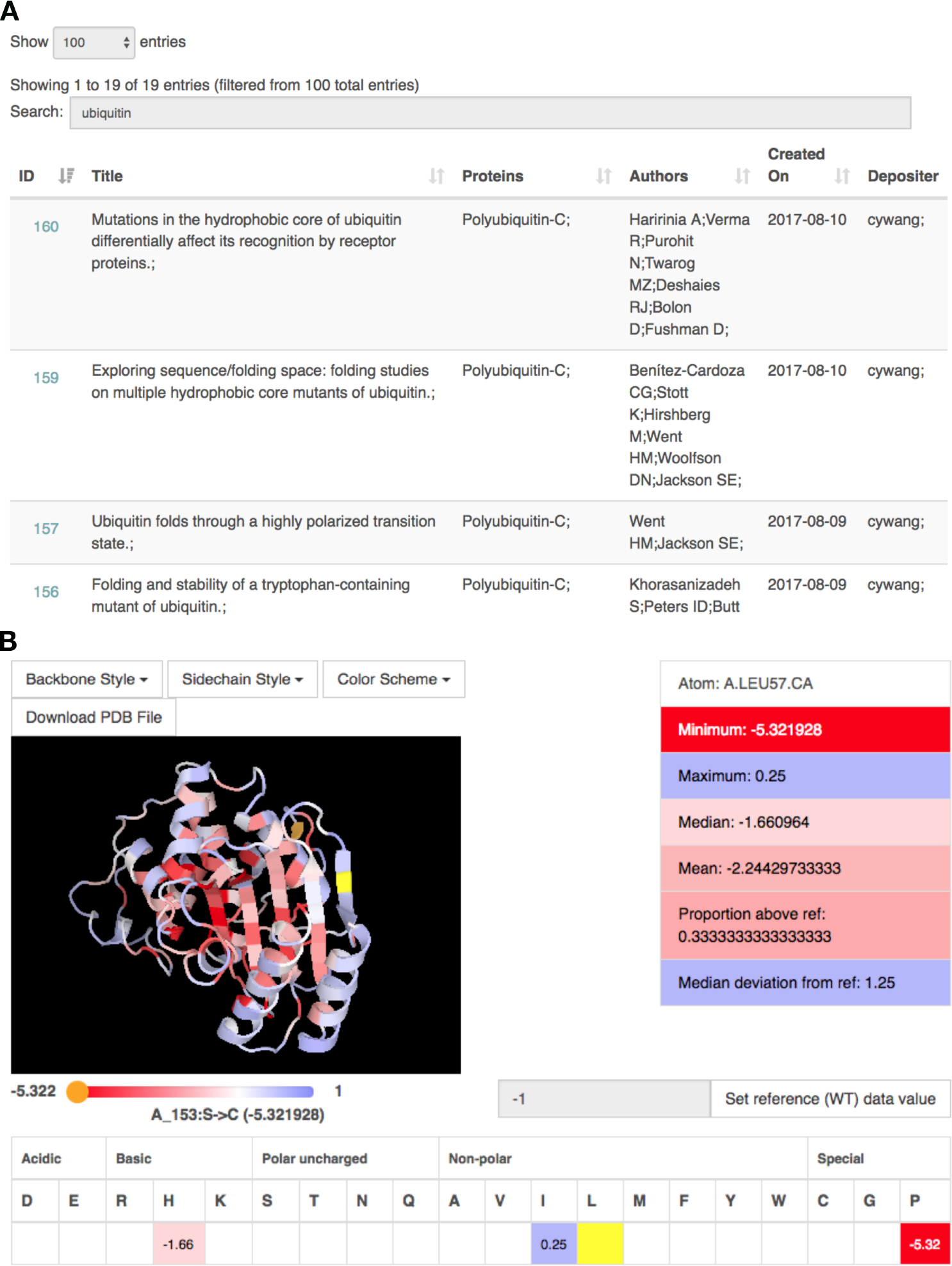
Screenshots of the web interface when using ProtaBank search and analysis tools. (A) A text-based search for “ubiquitin” returns a sortable table containing all studies with ubiquitin in the protein name or study title. Clicking on the study ID at the left brings up the analysis page for that study. (B) The analysis page for a study on β-lactamase27 includes a protein visualizer in which mutational results are mapped onto the protein structure according to the selected color scheme. Here, Leu57 is highlighted in yellow and the single mutant data for that residue is displayed in the tables below and to the right. We see that Leu57 was mutated to His, Ile, and Pro, resulting in scores of −1.66, 0.25, and −15.32, respectively (mean = −2.24); residue values are colored from red to white to blue depending on whether they are less than (red) or greater than (blue) the value of the reference at that position (white).

## Utility and Discussion

The following case studies demonstrate how ProtaBank search and analysis tools can aid in analyzing and interpreting PE data.

### Case study 1: Identify and compare data for a protein sequence

Before beginning any PE study, a review of existing literature on the protein of interest provides a useful reference point. Therefore, a simple but important application of ProtaBank is to identify and compare previously measured properties of a given sequence. Because ProtaBank stores the full sequence information for each mutant, a simple query on a specified protein sequence retrieves all the relevant data for that sequence, even if the starting sequences were different. In this case study, we use ProtaBank’s “Compare data for a sequence” tool to search for data on the wild-type sequence of the β1 domain of Streptococcal protein G (Gβ1):

MTYKLILNGKTLKGETTTEAVDAATAEKVFKQYANDNGVDGEWTYDDATKTFTVTE.

ProtaBank returns a sortable and searchable table listing all the data for the specified search sequence, including all the properties, assays, results, units, and titles of the associated studies. We can then search this data table for “Gibbs free energy” to just show the data in which Δ*G*s were measured. The Δ*G* search shows five experimentally measured values for Δ*G* of unfolding (Δ*G*_u_) from five studies,^29–33^ with values differing by up to 1.8 kcal/mol.

These differences could represent statistical variation in the measurement of this property. However, differences in assay techniques or conditions could also be responsible. ProtaBank provides links in the data table so that you can quickly view the details for each assay. A careful examination of assay details shows that an important difference between the assays was the pH used for denaturation; the temperature was 25°C for all the measurements except one (see Table I). Different techniques were also used (chemical vs. thermal denaturation), but these gave similar results when the temperature and pH were similar. These results suggest that the pH and/or temperature can have a notable effect on Δ*G*_u_. Thus, in order to make meaningful comparisons of engineered mutants relative to the wild type, it is clearly important to select the results with the most closely matched experimental conditions. We expect that by facilitating these types of comparisons, ProtaBank will provide context for the results in each study, reveal assay parameters that can impact the results, and enable an informed evaluation of results obtained under different assay conditions.

**Table I.** Assay Details Help Explain Differences in ΔG_u_ Results for wild-type Gβ1

| Study | Reference | $\Delta G_u$ (kcal/mol)<sup>a</sup> | Technique<sup>b</sup> | T (°C)<sup>c</sup> | pH |
| --- | --- | --- | --- | --- | --- |
| 57 | Choi and Mayo, 2006<sup>32</sup> | 5.9<sup>d</sup> | Thermal denaturation, circular dichroism | 25 | 5.5 |
| 61 | Gronenborn *et al.*, 1996<sup>29</sup> | 5.6 | GdmCl denaturation, fluorescence | 25 | 5.4 |
| 72 | Frank *et al.*, 1995<sup>30</sup> | 4.1 | Urea denaturation, fluorescence | 25 | 2.0 |
| 74 | Kuszewski *et al.*, 1994<sup>31</sup> | 4.8 | GdmCl denaturation, fluorescence | 5 | 4.0 |
| 171 | Davey *et al.*, 2017<sup>33</sup> | 4.1 | GdmCl denaturation, fluorescence | 25 | -<sup>e</sup> |

<sup>a</sup> $\Delta G_u$ , Gibbs free energy of unfolding; <sup>b</sup>GdmCl, guanidinium chloride; <sup>c</sup>T, temperature; <sup>d</sup>value was -5.9 for $\Delta G$ of folding; $\Delta G_u$ is therefore opposite in sign (5.9); <sup>e</sup>pH not reported.

For theoretical and computational scientists, ProtaBank provides another valuable service—easy access to data sets that can be used to benchmark, test, and improve predictive methods. For example, the experimental results provided in this case study could be used to test theoretical methods aimed at predicting the effect of pH and/or temperature on a protein’s stability based on how many ionizable side chains it contains.^34, 35^

### Case study 2: Identify and analyze data for closely related mutants of a protein sequence

Protein engineers are typically not only interested in the data reported for a given sequence, but in the data reported for closely related sequences. By comparing results between a sequence and its mutants, the effects of mutation at a given position can be determined. The knowledge gained can then be used to guide the selection of positions and mutations in future engineering efforts. In this case study, we use ProtaBank’s “Identify and analyze sequence mutations” tool to retrieve all the studies and assays containing data for sequences closely related to wild-type Gβ1. After entering the sequence in the search box, a BLAST search is performed to identify all related mutant sequences. The BLAST search currently identifies ~1.3 million sequences in ProtaBank that are closely related to wild-type Gβ1. Summary information is displayed in a mutant distribution heat map and a histogram showing the distribution of the number of mismatches (Fig. 4). The heat map [Fig. 4(A)] shows the number of sequences containing a mutation to a given amino acid at a given position; the wild-type residue for each position is shown in white. The heat map reveals that the T2Q mutation (chartreuse) occurs most frequently and that mutants at positions 39, 40, 41, and 54 (yellow-green) represent a large number of all the mutants identified. The T2Q mutation is often included in studies of Gβ1 to prevent cleavage of the N-terminal methionine by post-processing enzymes,^36^ and the preponderance of data for positions 39, 40, 41, and 54 is explained by a study that examined all possible combinations of mutations at these four positions, a total of 160,000 (20^4^) variants.^37^ The histogram [Fig. 4(B)] shows the number of sequences found at each mismatch level, where the number of mismatches is the number of mutations needed to go from a given mutant sequence to the search sequence. In this example, most of the sequences are two or three mutations away from the search sequence. These plots show information that can help users determine which positions and mutations have already been studied and which new ones they might want to consider in future work.

**Figure 4.**
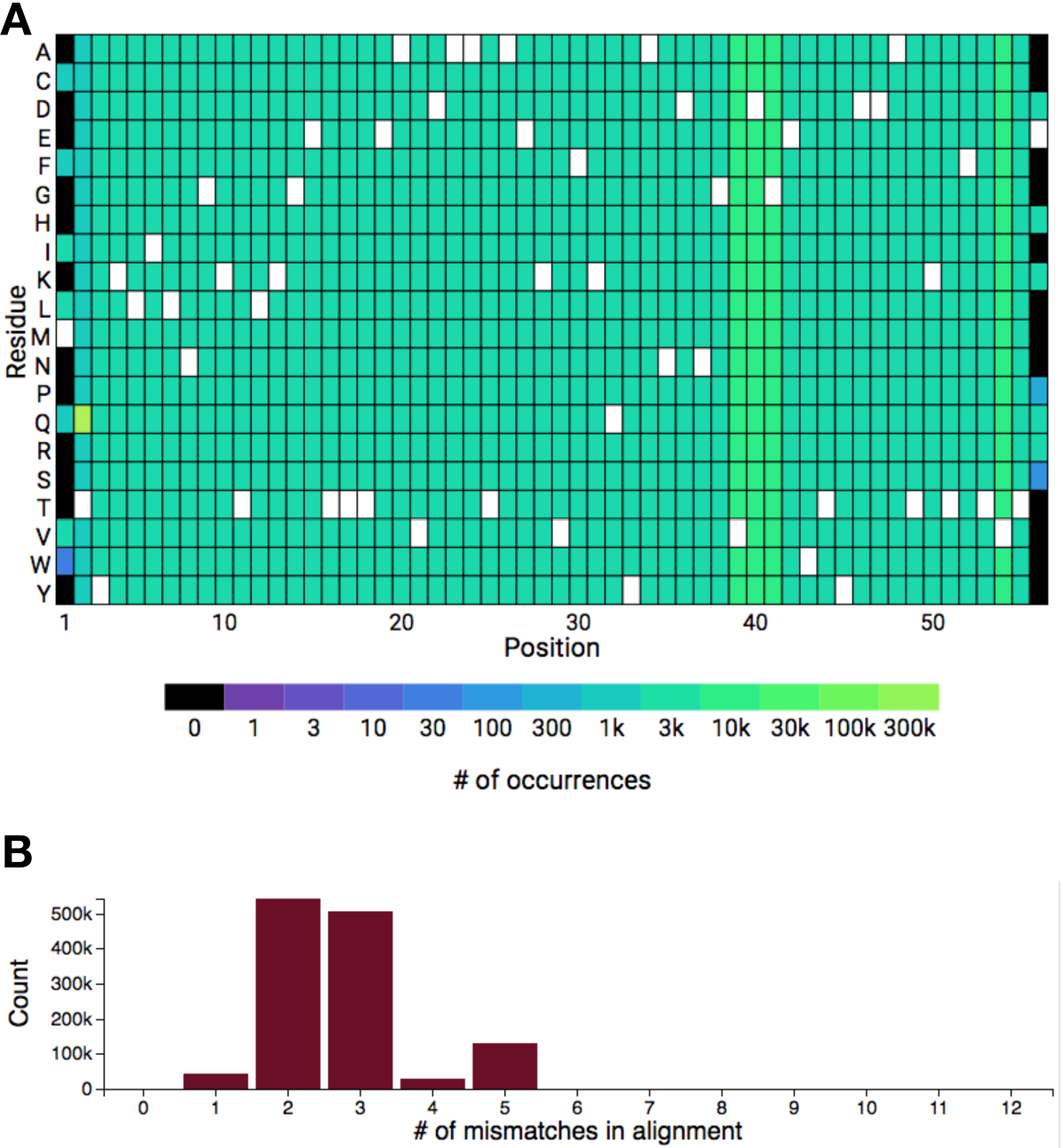
Identifying and analyzing closely related mutants of Gβ1 in ProtaBank. A BLAST search of the ProtaBank database finds ~1.3 million sequences that are closely related to wild-type Gβ1. (A) The heat map shows the frequency of each residue at each position. The wild-type residue is shown in white. (B) Histogram showing the number of sequences found at each mismatch level (Count), where the number of mismatches is the number of mutations needed to go from a given mutant sequence to the search sequence.

An “Assays by property” table is also displayed that lists all the assays containing data for a related mutant sequence, grouped by the protein property measured (Supporting Information, Fig. S2). For each property, the table lists all the individual assays, number of unique sequences, and total number of data points. Links to each of the assays provide quick access to assay details. Each of the data sets can be viewed via the # of data points link, which opens up a table displaying the results for that data set. This information can be downloaded as a CSV or Excel file.

### Case study 3: Determine the effects of mutations on protein properties and compare assay results

In this case study, we use additional features of ProtaBank’s “Identify and analyze sequence mutations” tool to perform further analyses on the closely related Gβ1 sequences retrieved in *Case study 2* above.

#### Plot one property vs. another

For any two measured properties, users can plot one property vs. another to show how these properties are correlated. ProtaBank automatically performs the unit conversions required to plot the data on the same set of axes. In Figure 5, we compare two measures of stability: *T*_m_ and the denaturant concentration at the midpoint of the unfolding transition (*C*_m_) for all the closely related sequences of Gβ1 retrieved in the BLAST search above (see *Case study 2*). A plot of all the Gβ1 mutant sequences for which both properties were measured (gray circles) shows a moderate correlation (r = 0.45) between these two properties, which could be explained by the fact that this comparison does not take differences in assay conditions into account. ProtaBank facilitates comparison of assay details by providing links to each of the assays listed in the “Assays by property” table (Supporting Information, Fig. S2). If *T*_m_ vs. *C*_m_ for the Gβ1 mutants is replotted using only data measured under similar assay conditions (e.g., pH, temperature, and denaturant), a very strong correlation is observed (r = 0.80) (Fig. 5, blue triangles).

**Figure 5.**
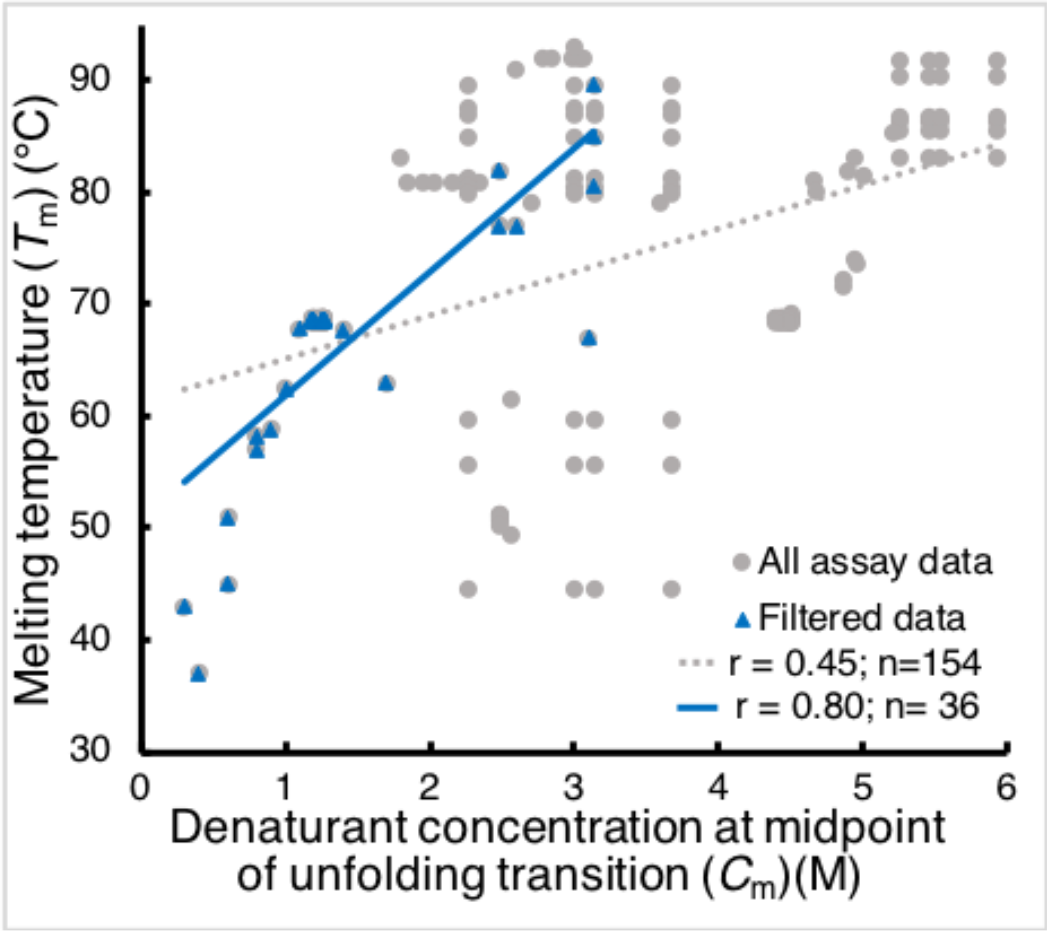
Plots of *C*_m_ vs. *T*_m_ for Gβ1 data. A plot of all Gβ1 mutant sequences for which both a *T*_m_ and *C*_m_ were measured (gray circles) gives a moderate correlation (r = 0.45, dotted gray line). If we only include data obtained under similar assay conditions (restricting *C*_m_ data to guanidinium chloride denaturation, pH 5-7, 20-30°C, and *T*_m_ data to pH 5-7, no denaturant added) (blue triangles), a much better correlation (r = 0.80, blue line) between these two measures of stability is observed.

#### Compare assay results

A recently published study by Olson *et al.*^38^ used mRNA display and deep mutational scanning to determine the fitness of all single and double mutants of Gβ1. The authors further calculated a ΔΔ*G* predictor (ΔΔ*G*_screen_), which used their fitness values to predict the Δ*G* change in protein stability upon single point mutations. The effectiveness of the predictor was evaluated by comparing the predicted results to experimentally obtained ΔΔ*G*s reported in the literature (ΔΔ*G*_literature_). ProtaBank provides a feature that allows this type of comparison to be done quickly and easily. The “Compare assay to others by mutation” feature allows all the input mutants for one assay to be searched for and compared to a given group of assays. ProtaBank automates the time-consuming task of manually identifying relevant literature results, converting the data to the same set of units, and displaying pertinent assay and background sequence information. All the results can then be further sorted and filtered by background sequence, mutation, or study. We used this feature to reproduce the comparison of ΔΔ*G*_screen_ to existing biochemical measurements of ΔΔ*G* as shown in the Olson *et al.* study.^38^ First, we did a “Compare assay to others by mutation” on the closely related sequences of wild-type Gβ1; this search identified hundreds of mutant sequence pairs in ProtaBank [Fig. 6(A)]. We then filtered these results to the set of 10 background sequences and single point mutants listed in the Olson *et al.* study [Fig. 6(B)]. Our filtered results match the data in their paper exactly except for one point—the mutant cited as I6L^29^ is actually a double mutant (I6L+T2Q) and was therefore excluded in our single mutant results. Note that ProtaBank identifies additional mutations not included in the Olson *et al.* data set and mutations with different background sequences (for a total of 90 unique background sequences), expanding the Olson *et al.* data set from 82 to 343 data points.

**Figure 6.**
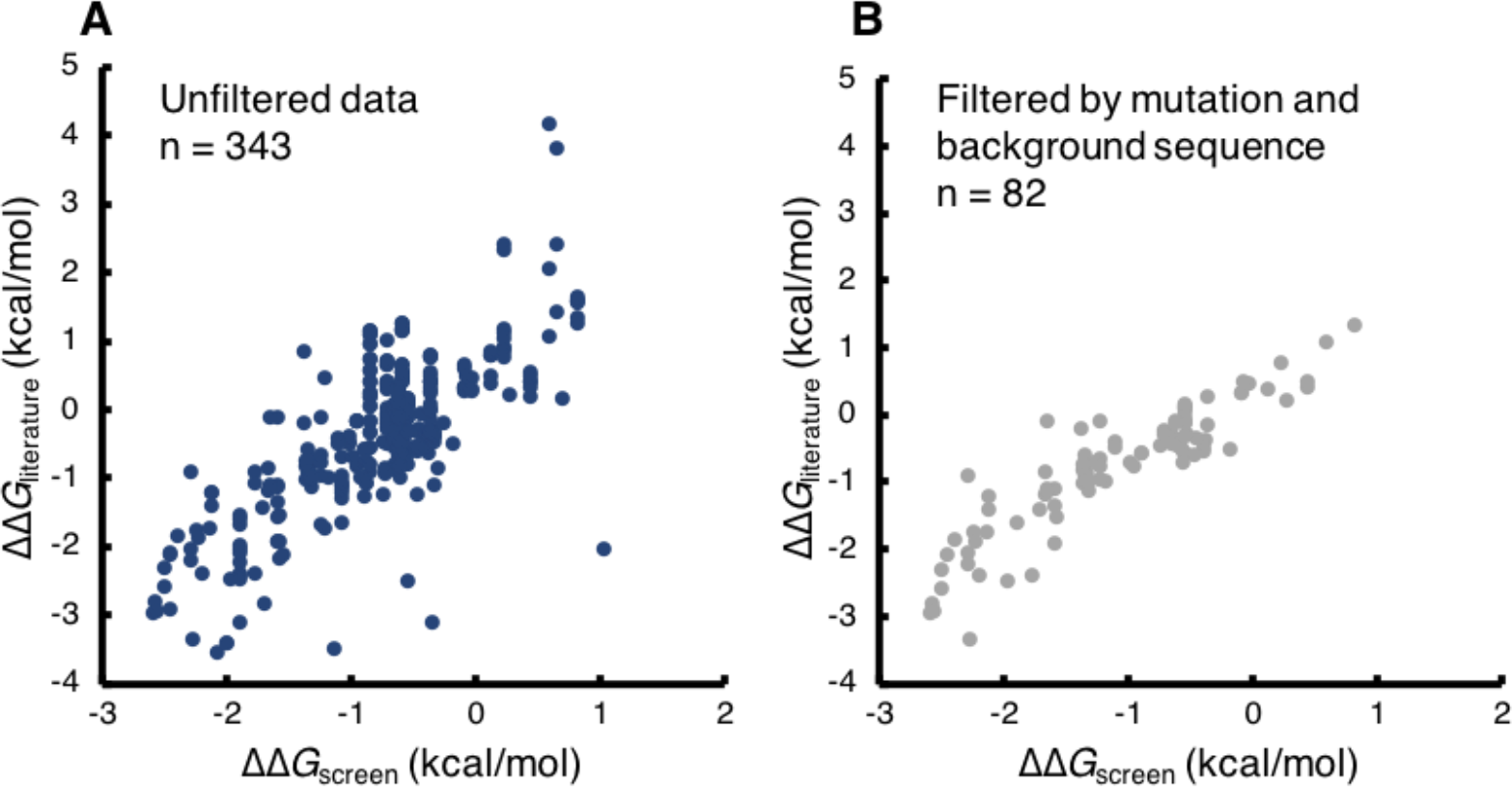
Comparing predicted with experimentally measured ΔΔ*G* values in ProtaBank. ProtaBank search tools were used to reproduce data from a study by Olson *et al.*^38^ in which Gβ1 fitness values were used to predict the change in stability upon point mutation. The ΔΔ*G* predictor values (ΔΔ*G*scree*n*) were plotted against experimental ΔΔ*G* values reported in the literature (ΔΔ*G*literature). (A) Unfiltered search of ProtaBank database identifies 343 mutant sequence pairs with both predicted and experimental ΔΔ*G* values. (B) Search filtered by the mutations and background sequences from the Olson *et al.* study yields 82 pairs, reproducing their data. Note that ProtaBank identifies ~260 additional data points.

This feature makes it easy to compare the results for the set of mutants in a given assay to those from any other group of assays (the properties measured can be the same or different). This allows one to see if new assay data is consistent with previously observed trends. It can also be used to identify protein properties that are well correlated with a particular assay.

### Case study 4: Visualize the relationship between mutations and protein structure

Often PE data can be better understood in the context of the protein’s 3D structure. In this case study, we look at experimental data from the Olson *et al.* Gβ1 study described above^38^ by mapping the effect of single mutations onto the crystal structure of the protein. By visualizing the data in this way, trends associated with structural features become more obvious than when viewed in a table or chart.

In the Olson *et al.* study, fitness values for every Gβ1 point mutant were determined by generating a DNA library encoding all single and double mutants and assessing relative binding affinity to IgG Fc. After a single round of affinity enrichment, the fitness of each variant was determined by the change in its frequency of occurrence (before vs. after enrichment). We can view this data in 3D with the protein visualizer, which is accessible via the study analysis page. We could map the fitness data onto the Gβ1 backbone using the median deviation from the wild-type value color scheme to help identify residue positions that are sensitive to mutation, as we did for the β-lactamase study in Figure 3(B). However, by just looking at the backbone image alone, it may not be immediately apparent why some residues are more sensitive to mutation than others.

Further analysis and visualization capabilities are therefore provided. ProtaBank allows you to save the data values from the selected color scheme in the occupancy column of the PDB file so that other modeling or visualization software can be used. In this case study, we used visual molecular dynamics (VMD)^43^ software to make the images shown in Figure 7. Two views of Gβ1 bound to the Fc domain (PDB ID: 1FCC)^44^ are displayed. On the left, the Gβ1 backbone is colored by median deviation from the wild-type value, with large deviations shown in blue, medium in white, and small to no deviations in red. On the right, the backbone is colored by proximity to the binding site: residues within 3.0 Å of the Fc domain are shown in blue, those between 3.0 and 3.5 Å from the Fc domain are shown in white, and those greater than 3.5 Å away are shown in red. Note that most of the residues near the binding site (right, blue or white) also show large median deviations from the wild-type value (left, blue to white). These results are understandable given that the study employed a selection assay based on Fc binding. The structural analysis thus helps explain why these residues are particularly sensitive to mutation and suggests that the observed sensitivity is likely due to disruption of the binding site rather than a destabilization of the Gβ1 fold.

**Figure 7.**
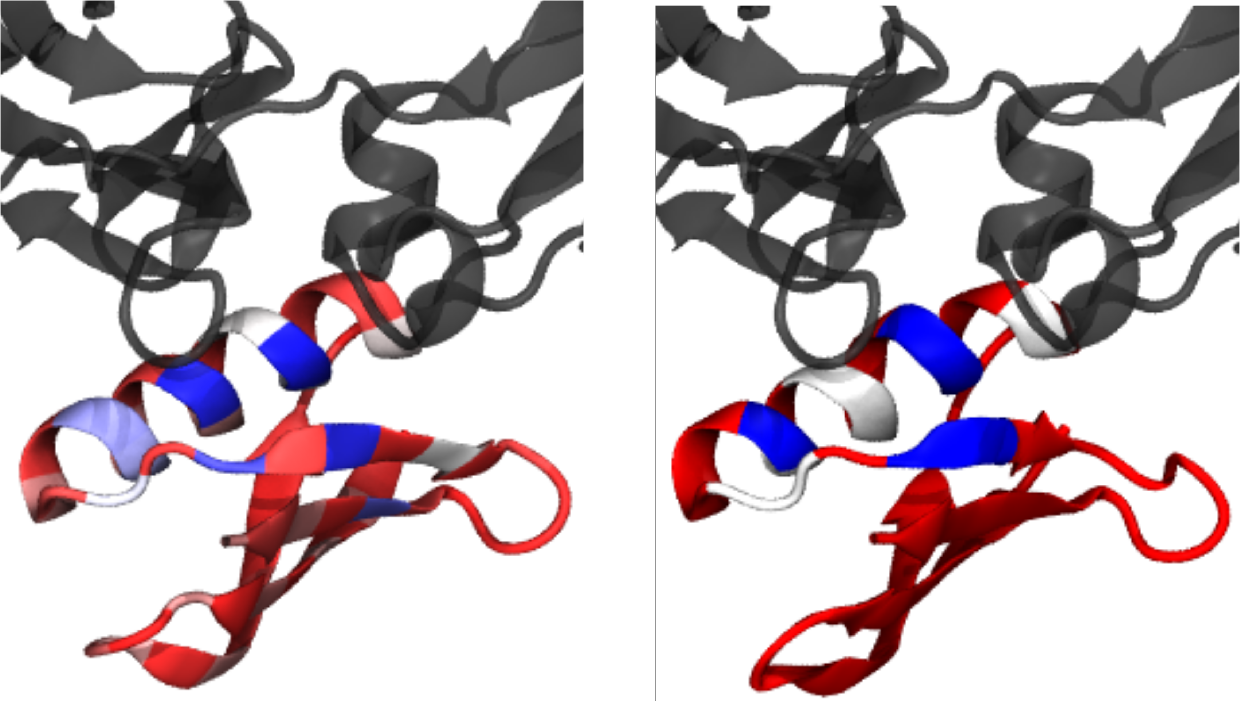
Comparing fitness and proximity to the binding site for GC1 point mutants. The ProtaBank visualizer was used to map the Olson *et al.*^38^ fitness data to the Gf1 structure and make the two images shown here. Gβ1 (red, white, and blue) is displayed bound to the Fc domain (gray) (PDB ID: 1FCC).^44^ On the left, the Gβ1 backbone is colored by median deviation from the wild-type value, going from blue to white to red, with large deviations in blue, medium in white, and small to no deviations in red. On the right, the backbone is colored by proximity to the binding partner: blue if within 3.0 Å of the Fc domain, white if between 3.0 and 3.5 Å and red if more than 3.5 Å away. The structural analysis shows that most of the Gβ1 residues near the binding interface are particularly sensitive to mutation.

## Concluding Remarks and Future Development

ProtaBank offers an easily accessible cloud-based modern database for PE data. It emphasizes the specification of detailed assay information and full protein sequences in an effort to ensure that all collected data is not just stored, but that data from diverse studies are comparable, searchable, and easily analyzed. ProtaBank has a convenient web interface to facilitate data entry for single studies and a REST API to allow for the upload of large data sets. By accepting data submissions directly from researchers, ProtaBank can incorporate the most recent results and be managed with fewer resources. Although this requires some effort on the part of the individual researcher, ProtaBank offers many benefits to submitters, including storing their data in an organized format on the cloud and allowing results to be searched and viewed by the scientific community, thereby increasing its impact.

In future development, we will expand ProtaBank’s analysis and data mining tools. The current analysis tools allow users to identify relevant data, find correlations between types of data, create plots and charts, and view results on the 3D structure. We have also started more advanced integration with protein structural data to allow for data selection and filtering on structural properties and to allow for computational predictions based on structural and sequence information. Future tools include incorporating computational methods to predict the effect of mutations on protein properties such as stability, binding, and activity.

ProtaBank will provide a central location and valuable entry point for researchers to store, retrieve, compare, and analyze PE data. It will make it easier for scientists to find previous results to guide their designs and provide valuable data sets that theoreticians can use as benchmarking cases in developing better predictive algorithms. We expect that ProtaBank will serve a pivotal role in centralizing PE data and leveraging the increasingly large amount of mutational data being generated. ProtaBank and its analysis tools will accelerate our ability to understand sequence-function relationships and greatly facilitate future protein design and engineering efforts.

## Supplementary Material

Supplementary material includes a table listing the protein properties included in ProtaBank, a figure showing a screenshot of the data entry page for a mutant library, and a figure showing results obtained from a sequence search.

## Acknowledgments

This work was supported by the National Institute of General Medical Sciences of the National Institutes of Health under Award Number R44GM117961. The content is solely the responsibility of the authors and does not necessarily represent the official views of the National Institutes of Health.

## Supporting Information

### ProtaBank: A repository for protein design and engineering data

Connie Y. Wang, Paul M. Chang, Marie L. Ary, Benjamin D. Allen, Roberto A. Chica, Stephen L. Mayo and Barry D. Olafson

**Table S1.** Protein properties included in ProtaBank

| Category | Properties |
| --- | --- |
| Activity | Activity, Catalytic efficiency ($k_{\text{cat}}/K_m$), Catalytic rate constant ($k_{\text{cat}}$), Count/Number, EC50, Energy, Enrichment, Epistasis, Fitness, IC50, Inhibition constant ($K_i$), Maximal rate ($V_{\text{max}}$), Michaelis constant ($K_m$), Relative activity, Specific activity |
| Binding | Association constant ($K_a$), Count/Number, Dissociation constant ($K_d$), ELISA, Energy, Enrichment, Enthalpy of binding ($\Delta H$), Entropy of binding ($\Delta S$), Epistasis, Fitness, Frequency of occurrence, Gibbs free energy of binding ($\Delta G$), Inhibition constant ($K_i$), Rate constant of association ($k_{\text{on}}$), Rate constant of dissociation ($k_{\text{off}}$) |
| Expression | Concentration, Energy, Optical density (OD), Enrichment, Frequency of occurrence, Minimum inhibitory concentration (MIC), Yield |
| Growth | Antimicrobial resistance, Energy, Enrichment, Frequency of occurrence, Optical density (OD) |
| Preclinical/Clinical | Bioavailability, EC50, Half-life ($t_{1/2}$), IC50, Immunogenicity, Toxicity |
| Solubility/Aggregation | Concentration, Energy, Fractional increase in solubility, Insoluble fraction, Oligomerization state, Soluble fraction |
| Specificity | Energy, Frequency of occurrence, Relative activity, Relative affinity, Relative $k_{\text{cat}}$, Relative $k_{\text{cat}}/K_m$, Relative $K_d$ |
| Spectral properties | Brightness, Emission wavelength ($\lambda_{\text{em}}$), Energy, Excitation wavelength ($\lambda_{\text{ex}}$), Extinction coefficient, Fluorescence intensity, Maturation half-time, Photobleaching half-time, pKa, Quantum yield |
| Stability/Folding | Constant pressure heat capacity of unfolding ($\Delta C_p$), Count/Number, Denaturant concentration at midpoint of unfolding transition ($C_m$), Energy, Enthalpy of unfolding ($\Delta H$), Entropy of unfolding ($\Delta S$), Equilibrium constant ($K$), Gibbs free energy of folding/unfolding ($\Delta G$), Melting temperature ($T_m$), β-Tanford value, Rate of folding ($k_F$), Rate of unfolding ($k_U$), Slope of chevron plot (m), Slope of the denaturant unfolding curve/cooperativity value (m), Temperature of maximum stability, Φ-value |

*Note:* Additional categories and/or properties can be specified by the user. Standard deviation or standard error data can be saved for any property.

**Figure S1.**
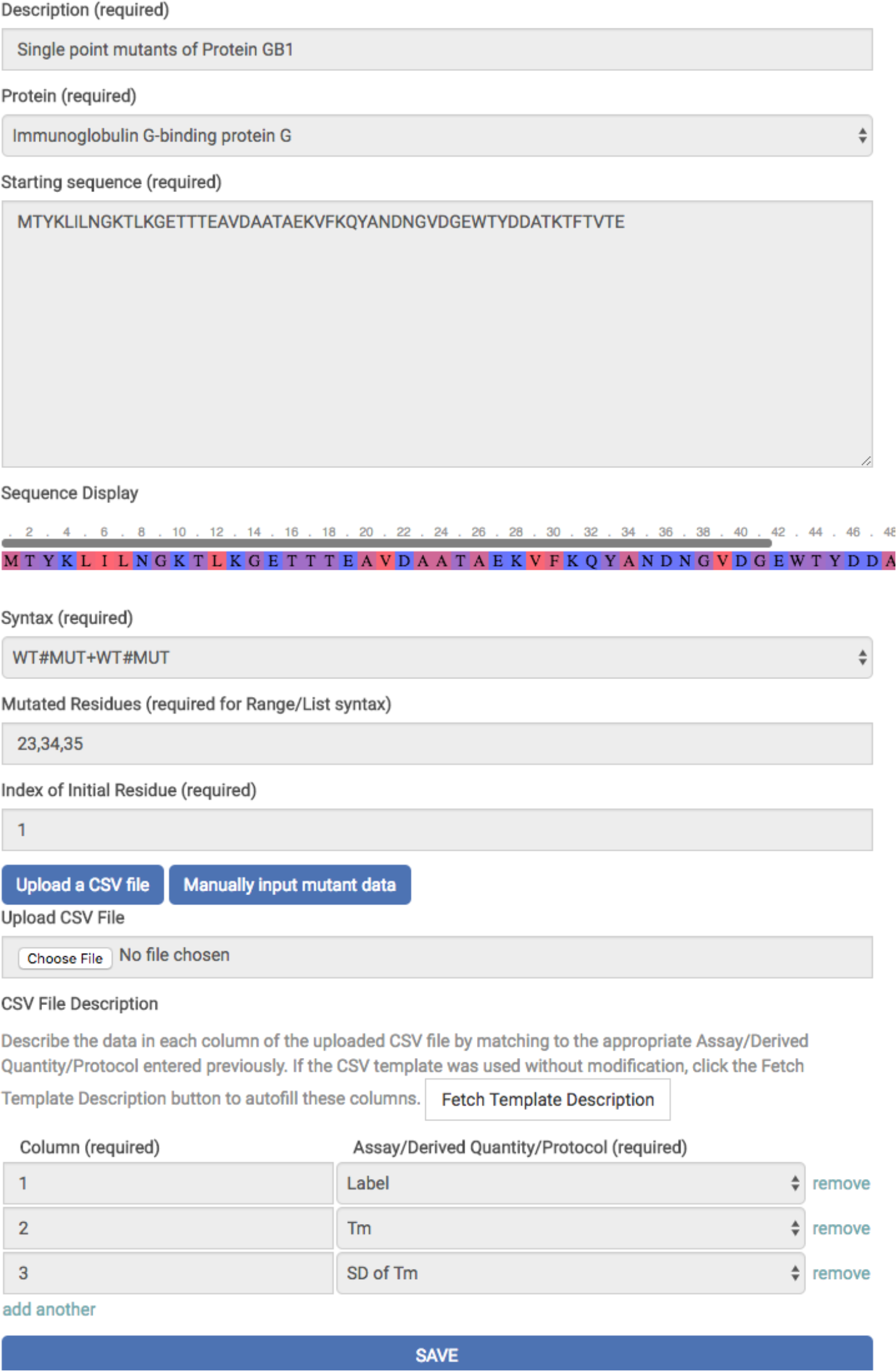
ProtaBank data deposition form for data in a mutant library. A short description of the data set, the protein that was mutated, and the starting sequence are specified, along with the format (syntax) used to describe the mutants. The mutational data can be uploaded from a CSV file or entered manually. This screenshot shows the web form for a CSV file upload: the data in each column of the CSV file is specified by selecting the appropriate Assay/Derived Quantity/Protocol for the property from the drop-down menu.

**Figure S2.**
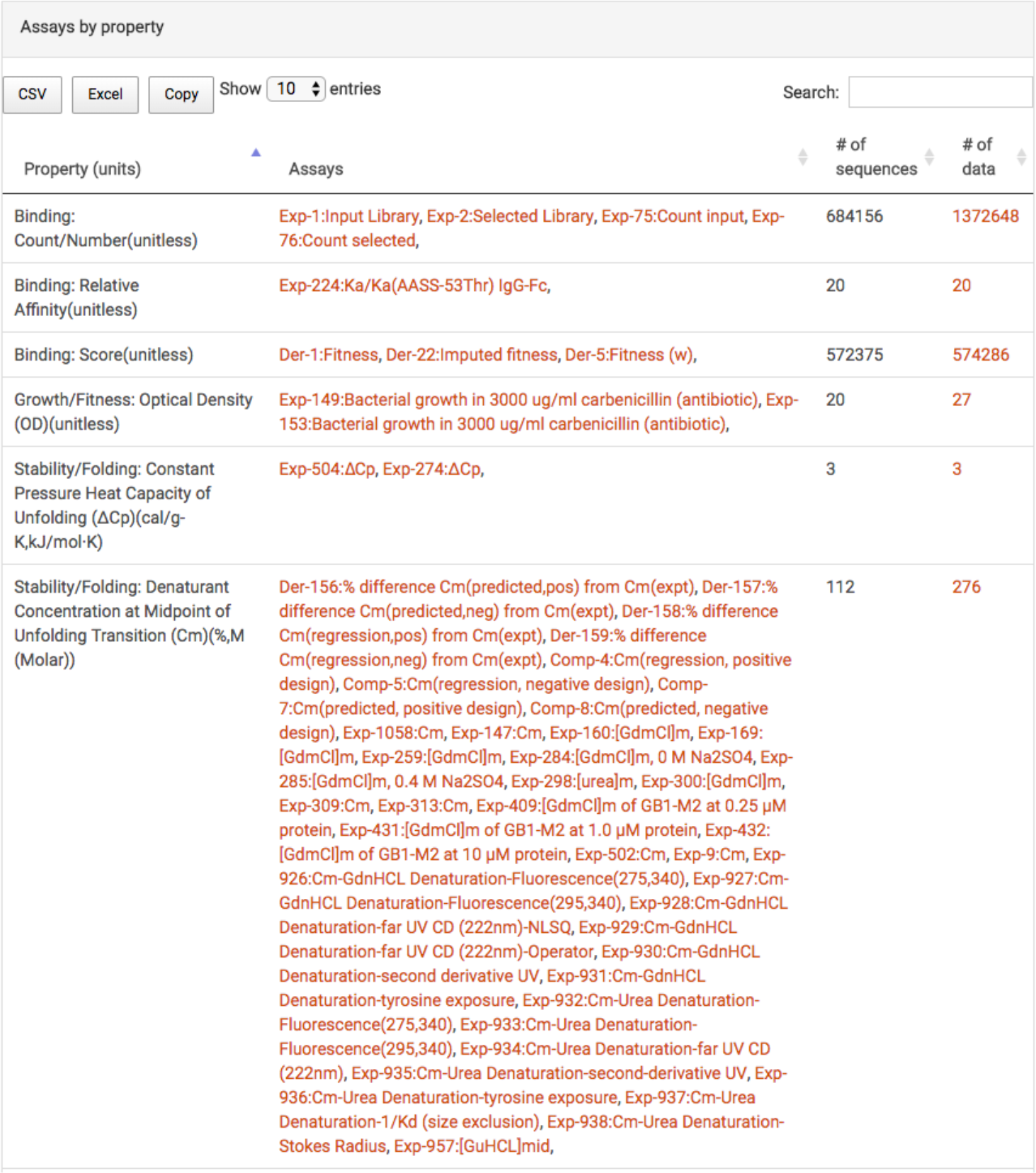
Assays by property table obtained using “Identify and analyze sequence mutations” tool to do BLAST search on wild-type Gβ1. All assays containing data for any of the closely related mutant sequences retrieved in the search are listed, grouped by the property measured; the data source for each assay is indicated with a prefix: Exp, experimental measurement; Comp, computational/simulated result; Der, derived from previously reported assay. The number of unique sequences and total number of data points are also given. Links (red) to each of the assays allow users to view assay details, and the “# of data” links bring up a table showing the results for that data set.

## References

1. Goodwin S, McPherson JD, McCombie WR (2016) Coming of age: ten years of next-generation sequencing technologies. Nat Rev Genet 17:333–351.

2. Romero PA, Tran TM, Abate AR (2015) Dissecting enzyme function with microfluidic-based deep mutational scanning. Proc Natl Acad Sci USA 112:7159–7164.

3. Chen B, Lim S, Kannan A, Alford SC, Sunden F, Herschlag D, Dimov IK, Baer TM, Cochran JR (2016) High-throughput analysis and protein engineering using microcapillary arrays. Nat Chem Biol 12:76–81.

4. Quan J, Saaem I, Tang N, Ma S, Negre N, Gong H, White KP, Tian J (2011) Parallel on-chip gene synthesis and application to optimization of protein expression. Nat Biotechnol 29:449–452.

5. Fowler DM, Araya CL, Fleishman SJ, Kellogg EH, Stephany JJ, Baker D, Fields S (2010) High-resolution mapping of protein sequence-function relationships. Nat Methods 7:741–746.

6. Hietpas RT, Jensen JD, Bolon DNA (2011) Experimental illumination of a fitness landscape. Proc Natl Acad Sci USA 108:7896–7901.

7. Whitehead TA, Chevalier A, Song Y, Dreyfus C, Fleishman SJ, De Mattos C, Myers CA, Kamisetty H, Blair P, Wilson IA, Baker D (2012) Optimization of affinity, specificity and function of designed influenza inhibitors using deep sequencing. Nat Biotechnol 30:543–548.

8. Fowler DM, Fields S (2014) Deep mutational scanning: a new style of protein science. Nat Methods 11:801–807.

9. Wrenbeck EE, Faber MS, Whitehead TA (2017) Deep sequencing methods for protein engineering and design. Curr Opin Struct Biol 45:36–44.

10. Rose PW, Prlić A, Altunkaya A, Bi C, Bradley AR, Christie CH, Costanzo LD, Duarte JM, Dutta S, Feng Z, Green RK, Goodsell DS, Hudson B, Kalro T, Lowe R, Peisach E, Randle C, Rose AS, Shao C, Tao Y-P, Valasatava Y, Voigt M, Westbrook JD, Woo J, Yang H, Young JY, Zardecki C, Berman HM, Burley SK (2017) The RCSB protein data bank: integrative view of protein, gene and 3D structural information. Nucleic Acids Res 45:D271–281.

11. Berman HM, Westbrook J, Feng Z, Gilliland G, Bhat TN, Weissig H, Shindyalov IN, Bourne PE (2000) The Protein Data Bank. Nucleic Acids Res 28:235–242.

12. Benson DA, Cavanaugh M, Clark K, Karsch-Mizrachi I, Ostell J, Pruitt KD, Sayers EW (2017) GenBank. Nucleic Acids Res 46:D41–D47.

13. Kumar MDS, Bava KA, Gromiha MM, Prabakaran P, Kitajima K, Uedaira H, Sarai A (2006) ProTherm and ProNIT: thermodynamic databases for proteins and protein-nucleic acid interactions. Nucleic Acids Res 34:D204–206.

14. The UniProt Consortium (2017) UniProt: the universal protein knowledgebase. Nucleic Acids Res 45:D158–169.

15. Placzek S, Schomburg I, Chang A, Jeske L, Ulbrich M, Tillack J, Schomburg D (2017) BRENDA in 2017: new perspectives and new tools in BRENDA. Nucleic Acids Res 45:D380–388.

16. Kawabata T, Ota M, Nishikawa K (1999) The protein mutant database. Nucleic Acids Res 27:355–357.

17. Moal IH, Fernández-Recio J (2012) SKEMPI: a Structural Kinetic and Energetic database of Mutant Protein Interactions and its use in empirical models. Bioinformatics 28:2600–2607.

18. Jemimah S, Yugandhar K, Michael Gromiha M (2017) PROXiMATE: a database of mutant protein-protein complex thermodynamics and kinetics. Bioinformatics 33:2787–2788.

19. Sirin S, Apgar JR, Bennett EM, Keating AE (2016) AB-Bind: Antibody binding mutational database for computational affinity predictions. Protein Sci 25:393–409.

20. NCBI Resource Coordinators (2017) Database Resources of the National Center for Biotechnology Information. Nucleic Acids Res 45:D12–D17.

21. Madden T. The BLAST Sequence Analysis Tool. In: Hoeppner M, Ostell J, Eds. (2013) The NCBI Handbook [Internet]. National Center for Biotechnology Information, Bethesda, MD, https://www.ncbi.nlm.nih.gov/books/NBK153387/.

22. Gekko K, Timasheff SN (1981) Mechanism of protein stabilization by glycerol: preferential hydration in glycerol-water mixtures. Biochemistry 20:4667–4676.

23. Scharnagl C, Reif M, Friedrich J (2005) Stability of proteins: temperature, pressure and the role of the solvent. Biochim Biophys Acta 1749:187–213.

24. Talley K, Alexov E (2010) On the pH-optimum of activity and stability of proteins. Proteins 78:2699–2706.

25. Kolesnikov N, Hastings E, Keays M, Melnichuk O, Tang YA, Williams E, Dylag M, Kurbatova N, Brandizi M, Burdett T, Megy K, Pilicheva E, Rustici G, Tikhonov A, Parkinson H, Petryszak R, Sarkans U, Brazma A (2015) ArrayExpress update–-simplifying data submissions. Nucleic Acids Res 43:D1113–1116.

26. Lee RYN, Howe KL, Harris TW, Arnaboldi V, Cain S, Chan J, Chen WJ, Davis P, Gao S, Grove C, Kishore R, Muller H-M, Nakamura C, Nuin P, Paulini M, Raciti D, Rodgers F, Russell M, Schindelman G, Tuli MA, Van Auken K, Wang Q, Williams G, Wright A, Yook K, Berriman M, Kersey P, Schedl T, Stein L, Sternberg PW (2017) WormBase 2017: molting into a new stage. Nucleic Acids Res 46:D869–D874.

27. Jacquier H, Birgy A, Le Nagard H, Mechulam Y, Schmitt E, Glodt J, Bercot B, Petit E, Poulain J, Barnaud G, Gros P-A, Tenaillon O (2013) Capturing the mutational landscape of the beta-lactamase TEM-1. Proc Natl Acad Sci USA 110:13067–13072.

28. Jelsch C, Mourey L, Masson JM, Samama JP (1993) Crystal structure of Escherichia coli TEM1 beta-lactamase at 1.8 A resolution. Proteins 16:364–383.

29. Gronenborn AM, Frank MK, Clore GM (1996) Core mutants of the immunoglobulin binding domain of streptococcal protein G: stability and structural integrity. FEBS Lett 398:312–316.

30. Frank MK, Clore GM, Gronenborn AM (1995) Structural and dynamic characterization of the urea denatured state of the immunoglobulin binding domain of streptococcal protein G by multidimensional heteronuclear NMR spectroscopy. Protein Sci 4:2605–2615.

31. Kuszewski J, Clore GM, Gronenborn AM (1994) Fast folding of a prototypic polypeptide: the immunoglobulin binding domain of streptococcal protein G. Protein Sci 3:1945–1952.

32. Choi EJ, Mayo SL (2006) Generation and analysis of proline mutants in protein G. Protein Eng Des Sel 19:285–289.

33. Davey JA, Damry AM, Goto NK, Chica RA (2017) Rational design of proteins that exchange on functional timescales. Nat Chem Biol 13:1280–1285.

34. Schaefer M, Sommer M, Karplus M (1997) pH-dependence of protein stability: absolute electrostatic free energy differences between conformations. J Phys Chem B 101:1663–1683.

35. Warwicker J (1999) Simplified methods for pKa and acid pH-dependent stability estimation in proteins: removing dielectric and counterion boundaries. Protein Sci 8:418–425.

36. Smith CK, Withka JM, Regan L (1994) A thermodynamic scale for the beta-sheet forming tendencies of the amino acids. Biochemistry 33:5510–5517.

37. Wu NC, Dai L, Olson CA, Lloyd-Smith JO, Sun R (2016) Adaptation in protein fitness landscapes is facilitated by indirect paths. eLife 5:e16965.

38. Olson CA, Wu NC, Sun R (2014) A comprehensive biophysical description of pairwise epistasis throughout an entire protein domain. Curr Biol 24:2643–2651.

39. Minor DL, Kim PS (1994) Context is a major determinant of beta-sheet propensity. Nature 371:264–267.

40. Malakauskas SM, Mayo SL (1998) Design, structure and stability of a hyperthermophilic protein variant. Nat Struct Biol 5:470–475.

41. Park SH, O’Neil KT, Roder H (1997) An early intermediate in the folding reaction of the B1 domain of protein G contains a native-like core. Biochemistry 36:14277–14283.

42. Cregut D, Civera C, Macias MJ, Wallon G, Serrano L (1999) A tale of two secondary structure elements: when a beta-hairpin becomes an alpha-helix. J Mol Biol 292:389–401.

43. Humphrey W, Dalke A, Schulten K (1996) VMD: visual molecular dynamics. J Mol Graph 14:33–38.

44. Sauer-Eriksson AE, Kleywegt GJ, Uhlén M, Jones TA (1995) Crystal structure of the C2 fragment of streptococcal protein G in complex with the Fc domain of human IgG. Structure 3:265–278.

